# Caveolae Impacts Cellular RNA Levels through Transcription and Translational Processes

**DOI:** 10.1101/2021.04.29.442036

**Authors:** Androniqi Qifti, Shravani Balaji, Suzanne Scarlata

## Abstract

Caveolae are membrane domains that provide mechanical strength to cells and localize signaling molecules. Caveolae are composed of caveolin-1 or −3 (Cav1/3) molecules that assemble into domains with the help of cavin-1. Besides organizing caveolae, cavin-1, also known as Polymerase I and Transcript Release Factor (PTRF), promotes ribosomal RNA transcription in the nucleus. Cell expression of Cav1 and cavin-1 are linked. Here, we find that deforming caveolae by subjecting cells to mild osmotic stress (300 to 150 mOsm), changes the levels of cellular proteins (GAPDH, Hsp90 and Ras) change only when Cav1/cavin-1 levels are reduced suggesting link between caveolae deformation and protein expression. We find that this link may be due to relocalization of cavin-1 from the plasma membrane to the nucleus upon caveolae deformation caused by osmotic stress. Cavin-1 relocalization is also seen when Cav1-Gαq contacts change upon stimulation with carbachol. Cav1 and cavin-1 levels have profound effects on the amount of cytosolic RNA and the size distribution of these RNAs that in turn impact the ability of cells to form stress granules and RNA-processing bodies (p-bodies) that protect mRNA when cells are subjected to environmental stress. Studies using a cavin-1 knock-out cell line show adaptive changes in cytosolic RNA levels but a reduced ability to form stress granules. Our studies show that caveolae, through release of cavin-1, communicates mechanical and chemical cues to the cell interior to impact transcriptional and translational processes.

## INTRODUCTION

Caveolae are flask-shaped membrane invaginations that can flatten to provide more membrane area and are implicated in mechano- and electric sensation, endocytosis, and vasodilation through modulating the NO pathway (Bernatchez et al., 2011; Buwa et al., 2020; Parton, 2018; Parton et al., 2018). In previous studies, our lab found that populations of Gαq and their receptors reside in caveolae domains and this localization is assisted by interactions between Gαq and caveolin molecules (Calizo and Scarlata, 2012; Sengupta et al., 2008). Activation of Gαq by hormones or neurotransmitters strengthens these interactions resulting in enhancement calcium signals. Deformation of caveolae by mild osmotic stress disrupts this stabilization and returns calcium signals to levels observed in the absence of caveolae (Guo et al., 2011; Guo et al., 2015; Yang and Scarlata, 2017). Additionally, when cells are subjected to either bi-directional static or oscillating mechanical stretch, calcium release through activation of Gαq/PLCβ is intact, but contacts between Gαq and caveolin are disrupted (Aisiku et al., 2011; Qifti et al., 2019a). In another series of studies, we found that activation of the Gαq/PLCβ pathway is linked to regulation of GAPDH protein production but not Hsp90 (see (Philip et al., 2012)).

Cavins are a family of four proteins that regulate the curvature of the caveolae membrane by anchoring caveolins to the cytoskeleton (for reviews see (Briand et al., 2011; Kovtun et al., 2015; Williams and Palmer, 2014)). The most abundantly expressed is cavin-1, also known as Polymerase 1 and Transcript Release Factor (PTRF) or cav-p60. Since its discovery in 1998, cavin-1 has been found to be a necessary component of caveolae formation, by mediating the sequestration of caveolin into immobile caveolae (see (Briand et al., 2011)). Several studies, including those here, strongly suggest that expression of cavin-1 and caveolin are interdependent; cav-1 knockout mice have nearly no cavin-1 expression, and cavin-1 knockout mice have diminished cav-1 expression (Liu and Pilch, 2008; Williams and Palmer, 2014). When fibroblasts are swelled by a 10-fold decrease in osmotic strength, cavin-1 is released from the plasma membrane as caveolae disassemble to provide more membrane area (Sinha et al., 2011).

Before cavin-1 was identified as a structural adaptor for caveolae, it was recognized for its role in modulating cellular transcriptional activity (Jansa et al., 1998). Cavin-1/PTRF promotes ribosomal DNA transcription by binding to the 3’ pre-RNA, allowing the release of pre-RNA and Polymerase I from the transcription complex (Liu and Pilch, 2016a). Cavin-1 not only plays a role in transcript release, but it increases the overall rate of transcription in a concentration-dependent manner. In adipocytes, insulin stimulation causes phosphorylation of cavin-1 promoting its translocation from caveolae to the nucleus (Liu and Pilch, 2016a) suggesting a role in signal transduction. The importance of cavin-1 expression is seen in knock-out mice that show diverse abnormalities consistent with impaired ribosome biogenesis including abnormal growth failure, loss in fat, resistance to obesity, impaired exercise ability, muscle hypertrophy, altered cardiac and lung function (see (Liu, 2020)). Pertinent for this study are reports suggesting that increased cavin-1 expression promotes cellular stress responses to toxic agents which many be traced to binding to p53 in the cytosol (Bai et al., 2011).

The connection between cavin-1’s ability to regulate caveolae structure and promote ribosomal RNA suggests that any mechanism that destabilizes cavin-1’s interactions with caveolin including caveolae deformation, would impact transcription and transitional processes through cavin-1. Here, we show that changes in environmental conditions, such as mild osmotic stress, addition of neurotransmitter, or exposure toxins drives cavin-1 from the plasma membrane to impact RNA transcription and two processes that regulate protein translation (i.e. stress granules and p-body formation). Our results suggest that Cav1/3/cavin-1 acts as sensor to inform the cell interior of environmental stress.

## RESULTS

### Expression of Cav1 alters protein expression in response to stress

Our previous studies showed that Gαq/caveolin interactions link mechanical and calcium signals (Sengupta et al., 2008), and additionally that the Gαq/PLCβ pathway can influence the regulation of GAPDH but not Hsp90 (Philip et al., 2012). We first tested whether caveolae expression would affect regulation of GAPDH through its stabilization of Gαq.

Using rat aortic smooth muscle (A10) cells, we quantified the production of three distinct cell proteins, GAPDH along with Hsp90 and Ras for comparison, when Cav1, Gαq and PLCβ were downregulated. We first found that reducing Cav1 changes the level of actin and appear to reduce the level of other cellular proteins (see below) making direct impact of Cav1 levels difficult to assess. We therefore took a more indirect approach. Because caveolae provides mechanical strength to cells, we subjected cells to mild osmotic stress that will deform caveolae and eliminate its stabilization of the Gαq/PLCβ pathways (Guo et al., 2015). We reduced the osmolarity of the media from 300 to 150 mOsm in control cells and cells treated with siRNA(Cav1) and quantified changes in GAPDH, Hsp90 and Ras levels (**Fig. 1**, *Supplemental 1*). Differences between the two cell groups were not immediate but appeared ~12 hours after continuous osmotic stress suggesting that Cav1 impacts slower transcription and/or translation processes rather than more rapid degradation or other down-regulation mechanisms. In contrast to Cav1, down-regulating PLCβ1 or Gαq had little or no effect arguing that the Gαq/PLCβ pathway is not the primary factor underlying changes in protein production. Thus, Cav1 levels impact the ability of cells to produce proteins under hypo-osmotic stress conditions.

**Fig. 1.**
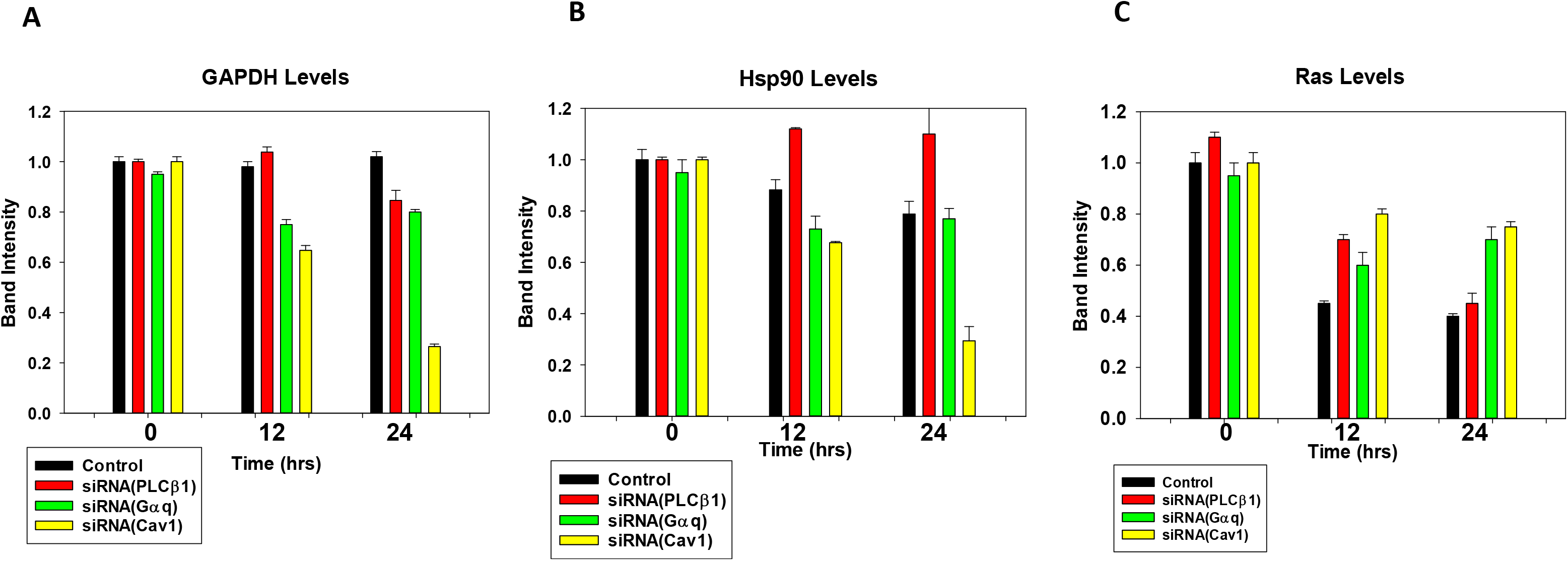
Down-regulation of Caveolin 1 impacts protein levels. Changes in proteins levels of **A-** GAPDH, **B-** Hsp90, **C-** Ras in rat aortic smooth muscle (A10) in control (black), siRNA PLC (red), siRNA Gαq (green) and siRNA Caveolin 1 (yellow) cells where n=3. An example western blot is shown in *Supplemental* **S1**.

### Cavin-1 shifts its cellular localization under osmotic stress

Expression of Cav1 and cavin-1 are interdependent ((Liu and Pilch, 2008) and *Supplemental 2*) and so we tested whether the protein changes observed in **Fig.2** might be due to cavin-1 levels since cavin-1 promotes transcription of ribosomal RNA (Jansa et al., 1998). While previous studies found high osmotic stress disassembles caveolae releasing cavin-1 from the plasma membrane (Sinha et al., 2011), it is unclear whether release will occur at the mild, ore physiological stress conditions used here which deform but not disassemble caveolae (Yang and Scarlata, 2017).

Cavin-1 relocalization studies were carried out in intact Wistar Kyoto smooth muscle cells (WKO-3M22) transfected with eGFP-cavin-1 and measuring shifts in the cellular distribution of fluorescence intensity with hypo-osmotic stress by confocal and total internal reflection (TIRF) microscopy. Under basal conditions, we find that eGFP-cavin-1 localizes mainly on the plasma membrane with small amounts in the cytoplasm and nucleus (**Fig. 2A**). Because the additional, transfected protein would increase total cellular cavin-1 and effect its localization, we repeated these studies by first down-regulating cavin-1 before transfecting with GFP-cavin-1. In this case, we find the intensity is slightly higher on the plasma membrane as compared to the nucleus, and none was seen in the cytosol suggesting that endogenous cavin-1 is mainly distributed between these two compartments.

**Fig. 2.**
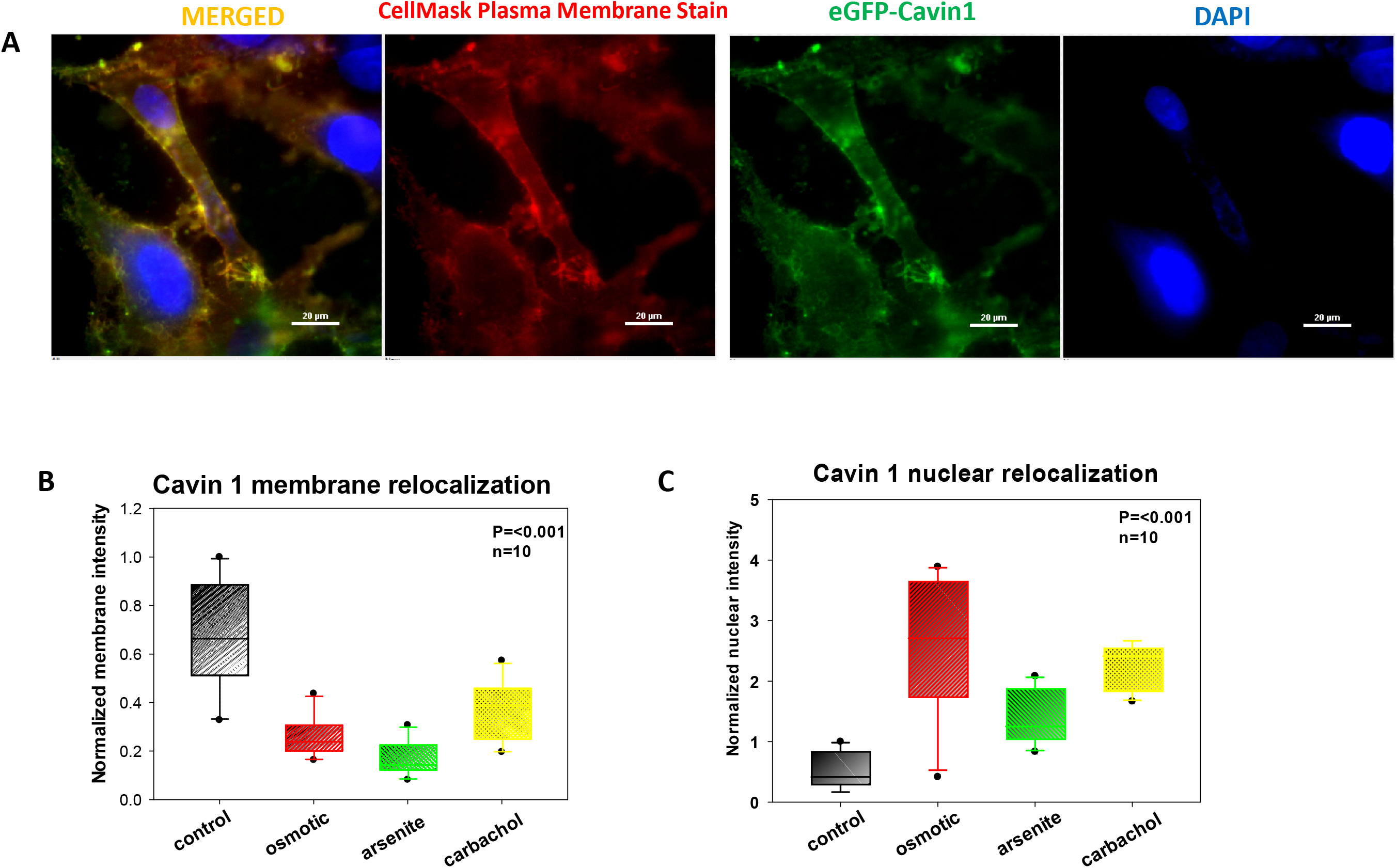
Cavin-1 relocalization in WKO-3M22 cells. **A-** Sample fluorescence images of fixed WKO-3M22 cells transfected with eGFP-Cavin1 (green) and stained with CellMask Deep Red Plasma Membrane stain (red) and DAPI (blue) under control conditions as obtained using Total Internal Reflection Fluoresence Microscopy (TIRF) with 60X magnification. Scale bars are 20 μm long. **B,C-** Cell localization of eGFP-cavin-1 was assessed in subjected to hypo-osmotic (red), arsenite (green) and Gαq stimulation with carbachol (yellow). eGFP-cavin-1 localization was determined by measuring the pixel intensities to coordinates to the plasma membrane, cytosolic and nucleus compartments based on the location of DAPI and the CellMask plasma membrane stain. Changes in membrane (**B**) and nuclear localization (**C**) in wild type cells. **D,E-** Sample fluorescence images of fixed WKO-3M22 cells transfected with eGFP-Cavin1 (green) and stained with CellMask Deep Red Plasma Membrane stain (red) and DAPI (blue) under hypo-osmotic conditions as obtained by (**D**) confocal microscopy with 40X magnification and (**E**) TIRF images with 60X magnification. Scale bars are 10 μm long. **F-H-** Cell localization of eGFP-cavin-1 in cells treated with siRNA(Cav1). Changes in localization in the nucleus (**F**), cytosol (**G**) and membrane compartments (**H)** were observed. All measurements are an average of 3 independent experiments that sampled 10 cells, where SD is shown and where the p values was determined using ANOVA.

**Fig. 2.**
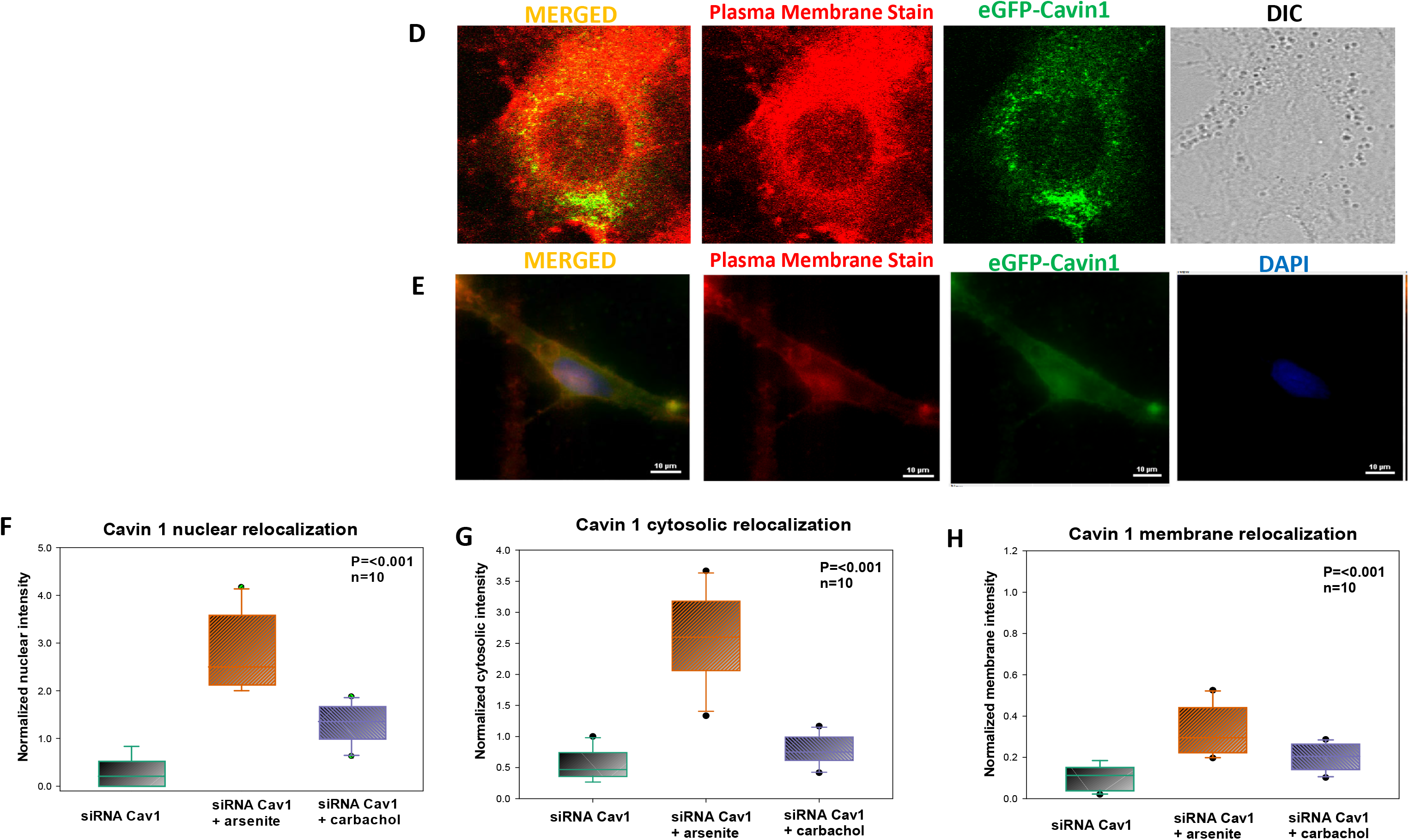
Cavin 1 relocalization in WKO-3M22 cells.

When we subjected wild type eGFP-cavin-1 cells to mild hypo-osmotic stress, a large portion of the plasma membrane GFP-cavin-1 population shifts to the cytosol and the nucleus (**Fig. 2B-2E**). To support the idea that cavin-1 moves off the plasma membrane when caveolae domains are disrupted, we down-regulated Cav-1 and found that little cavin-1 is associated with the plasma membrane under these conditions (**Fig. 2F-2H)**. Taken together, these data show that deformation, and not disassembly of caveolae, has the potential to promote nuclear localization of cavin-1 to impact transcription.

We also tested whether other stress conditions impact cavin-1 localization. Because Cav1/ Gαq interactions strengthen when is activated, we determined whether Gαq stimulation would in turn change cavin-1/Cav1 interactions and cavin-1 localization. To this end, we find that addition of carbachol to activate Gαq promotes relocalization of GFP-cavin-1 to the nucleus (**Fig.2**). We also found that treating cells with arsenite, which promotes a p53 response and is correlated to cavin-1 activity, also promotes cavin-1 nuclear localization, but to a lesser extent compared to Gαq activation (Kurki et al., 2003).

The data in **Fig.2** show that several stress conditions promote movement of cavin-1 from the plasma membrane to the nucleus, which would be expected to increase protein production based on cavin-1’s nuclear function. However, this is not seen in **Fig.1** where similar or lower levels of the protein, normalized for actin and changes in total protein content, were observed. Therefore, we investigated other mechanism through which Cav1 would influence protein levels.

### Transient down-regulation of Cav1 or cavin-1 affects the amount of cytosolic RNA and the size distribution

In the studies above, we casually observed that down-regulating Cav1 or cavin-1 results in increased cell death by 40% and 52%, respectively, as estimated by comparing the number of transfected to control cells in 3-5 culture dishes. We postulated that this increased mortality is due decreased transcription due to loss of cavin-1. To this end, we measured changes in the levels of cytosolic RNA (**Fig. 3A**). These studies were done by isolating the cytosolic fractions of WKO cells, removing the nuclear and lipid components, and isolating the RNA after protein digestion and lipid solubilization (see *Methods*). We find that down-regulating Cav1 by 56% and cavin-1 by 46%, as estimated by western blotting in WKO-3M22 cells (*Supplemental 2A*), resulted in a 5-fold reduction of cytosolic RNA while down-regulating cavin-1 by 70% and Cav1 by 82% resulted in a ~13 fold reduction (**Fig. 3A**). Notably, subjecting cells to osmotic strength also reduced cytosolic RNA levels even though more cavin-1 was nuclear (**Fig. 3C**)

**Fig. 3.**
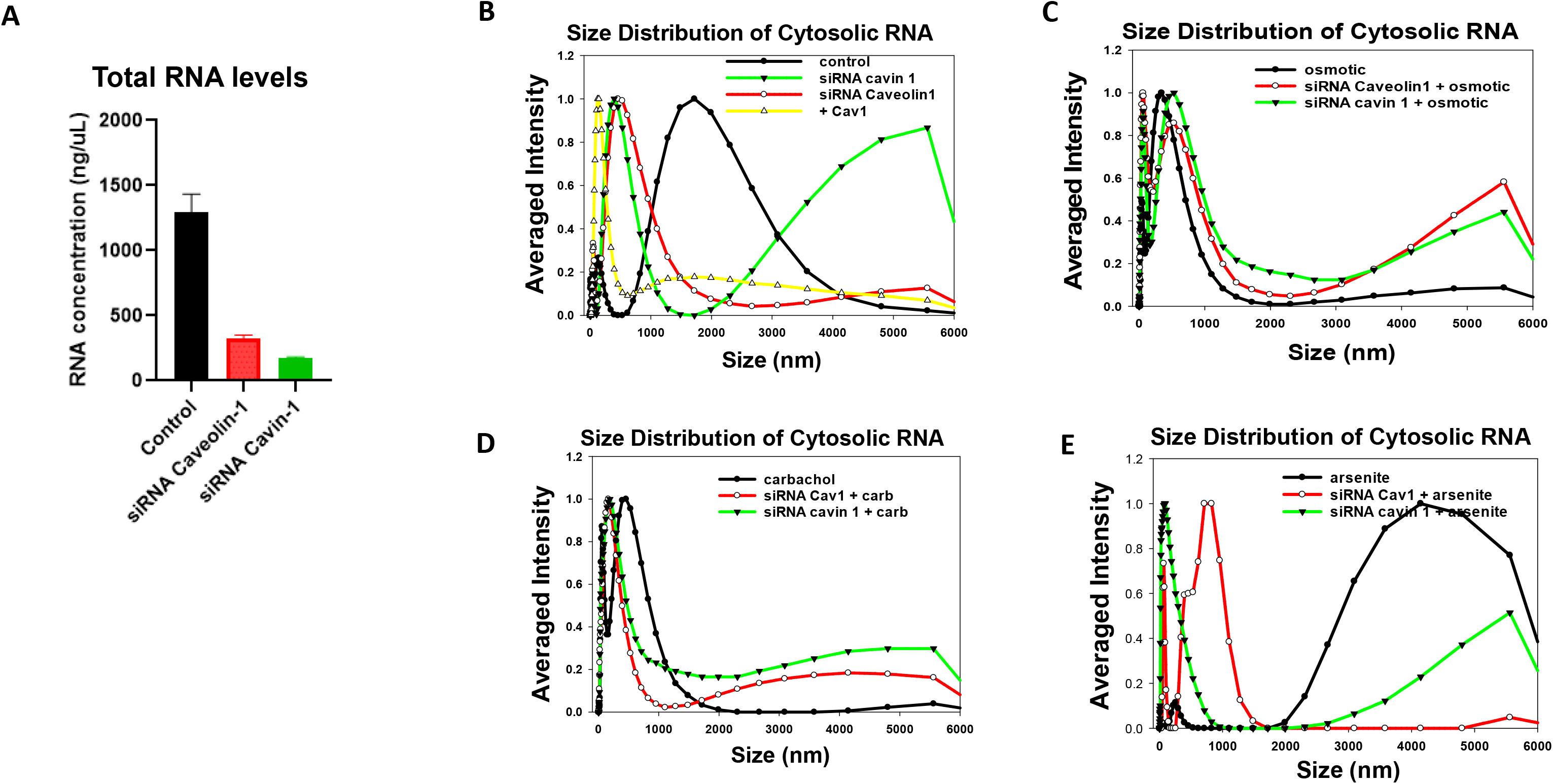
Characterization of RNA in WKO-3M22 cells. **A-** RNA from WKO-3M22 cells treated with siRNA(Cav1) (red) and siRNA(cavin-1) (green) was extracted and quantified (see methods). **B-** Normalized DLS spectra showing the size distributions of cytosolic RNA isolated from WKO-3M22 cells under control conditions (black), siRNA(Cav1) treated cells (red), cavin-1 overexpression (yellow), and siRNA(cavin-1) (green) down-regulation. **C-** Similar study as in (**B**) for cells subjected to hypo-osmotic stress (150 mOsm 5min), and **D-** cells subjected to Gαq stimulation with carbachol treatment (5μM, 10min). **E-** arsenite treatment (0.5mM, 10 min). Each sample was scanned 3 times with 10 minutes per run. The number of independent samples were 6 per condition.

We characterized relative size distributions of the cytosolic RNA populations by dynamic light scattering (DLS). We find that the transcripts effected by Cav1 down-regulation are smaller in size suggesting a higher level of processing. The RNA seen when cavin-1 is down-regulated is very large suggesting reduced processing, and the relative amount of the larger sized population decreases with the application of stress (**Fig. 3B-E**). The loss in RNA with Cav1/cavin-1 is consistent with reduced proliferation due to reduced transcription, and the shifts in the sizes of cytosolic RNA suggests that Cav1/cavin-1 levels impact processes related to RNA processing. We note that the percentage of rRNA to cytosolic RNA levels is reduced when Cav1 or cavin-1 s down-regulated as indicated by weaker contribution of the 18S and 28S ribosomal RNA bands (*Supplemental 2C*).

### Transient loss of Cav1/cavin-1 impacts stress responses

The changes in size distributions in **Fig. 3** suggest that Cav1/cavin-1 levels or stress conditions impact cytosolic RNA processing. To better understand these changes, we looked at the two major mechanisms used by cells to handle cytosolic RNA: processing bodies (p-bodies) and stress granules. P-bodies are associated with RNA degradation as well as RNA storage (Luo et al., 2018), and stress granules are halted ribosomal RNA complex that protect mRNA during stress conditions, such as the hypo-osmotic conditions used here (Anderson and Kedersha, 2006). Stress granules and p-bodies have an overlapping storage function and both contain Ago2 (Sen and Blau, 2005) which either stalls mRNA translation or degrades mRNA depending on the amount of base pairing between the Ago2-bound miR and the mRNA. Since both stress granules and p-bodies require RNA, their assembly will be impacted by the reduced RNA levels accompanying cavin-1 down-regulation.

We first tested whether Cav1/cavin-1 down-regulation changes the number and size of stress granules under basal conditions. These studies were carried out by immunostaining WKO-3M22 cells treated with either control or Cav-1 siRNA and immunostaining for the stress granule marker, polyadenylate binding protein (cytosolic) −1 (PABPC1). The size and area of PABPC1 particles were then quantified from high resolution fluorescence confocal images (see Methods). In **Fig 4A**, we show a comparison of control and siRNA (Cav1) in cells under basal conditions.

**Fig. 4.**
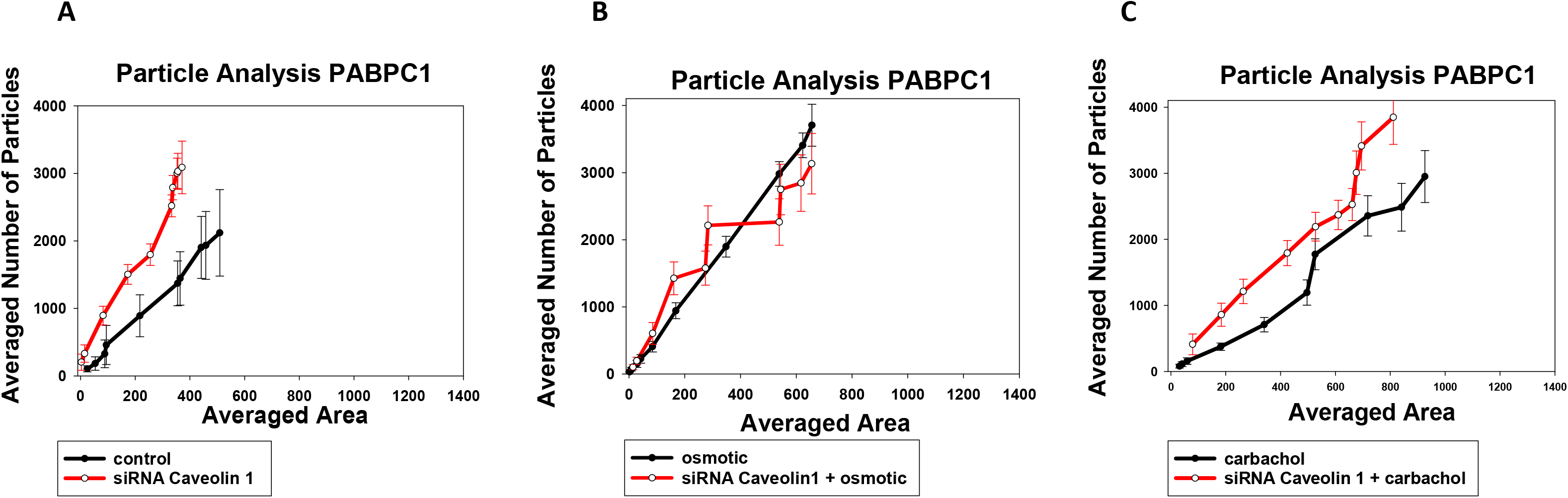
Caveolin 1 effects Stress Granule formation as seen by PABPC1 in WKO-3M22 Cells. The size and number of particles associated with monoclonal anti-PABPC1 in the cytosol of fixed and immunostained WKO-3M22 cells was measured on a 100x objective and analyzed using Image J (see methods). Comparison of cells with siRNA (Cav1) to mock-transfected under basal conditions (**A**), hypo-osmotic stress (150 mOsm, 5 minutes) (**B**), Stimulation of Gαq by treatment with 5 μM carbachol (**C**). All measurements are an average of 3 independent experiments that sampled 10 cells, where SD shown and where the p values was determined using ANOVA.

We find that loss of Cav1/cavin-1 results in more numerous but smaller particles consistent with smaller RNAs seen by DLS.

We then determined whether reducing Cav1/cavin-1 levels change the ability of cells to form PABPC1 particles (i.e. stress granules) under hypo-osmotic stress or carbachol stimulation (**Fig. 4B-C**). We find that reduced Cav1/cavin-1 levels did not affect particle formation in cells subjected to osmotic stress and produced a small increase in the number of fewer particles in cells stimulated with carbachol. These results suggest that the reduced protein production, such as the lower levels of GAPDH and Hsp90 seen in **Fig. 1** are not due to confinement in large stress granules.

The reduction in cytosolic RNA may result in smaller stress granules that might be missed in immunostaining studies. In a second series of studies, we monitored stress granule formation in live cells using the stress granule assembly protein G3BP1 tagged with GFP. This construct allowed us to monitor small aggregates by a fluorescence correlation method (N&B) which senses small aggregates. N&B is a fluorescence method which follows the diffusion of a fluorophore in an image series, and compares the number of photons associated with this diffusion to a monomeric control, such as free eGFP (Digman et al., 2008).

In **Fig 5**, we show control cells transfected with eGFP-G3BP1 and subjected to various stress conditions. The amount of aggregation with each stress condition correlated with cavin-1 relocalization seen in **Fig.2**; carbachol stimulation showed the highest amount of aggregation, followed by osmotic stress and then arsenite. When Cav1/cavin-1 was down-regulated, no changes in G3BP1 aggregation were observed except for osmotic stress which showed a small, residual change which might be due to residual Cav1. These results suggest that lowering Cav1/cavin-1 levels make cells less able to adopt mechanisms that protect them from environmental stress.

**Fig. 5-I.**
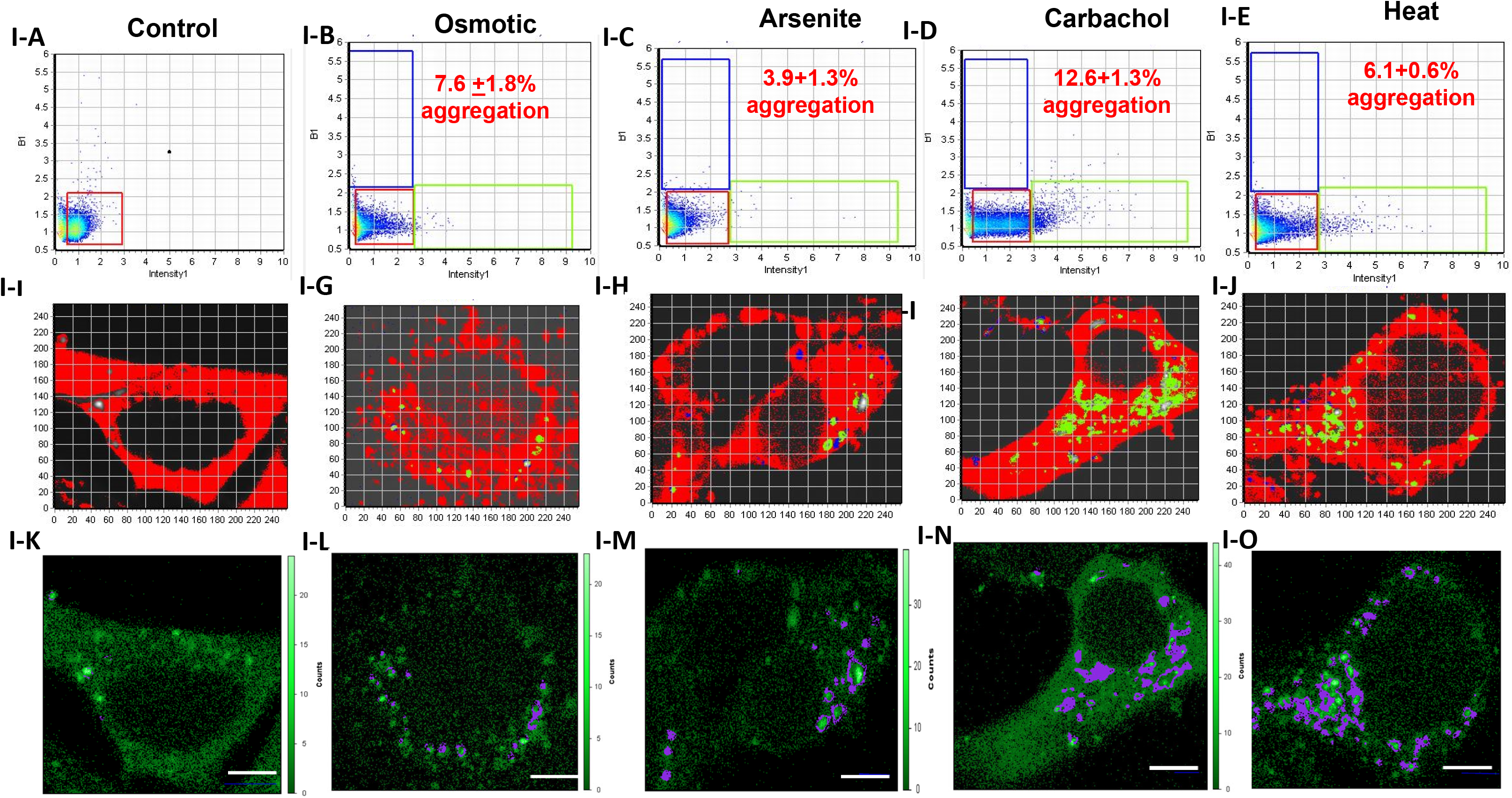
N&B analysis of eGFP-G3BP1 aggregation in WKO-3M22 cells. ***I-*** The top panels (**I-A – I-E**) show graphs of the brightness versus intensity with the pixels of the colored boxes corresponding to the specific regions in the cells (**I-F – I-J**) using SIM-FCS 4 software. The bottom panels show the corresponding fluorescence microscopy images in ISS (**I-K – I-O**). The red box corresponds to monomeric eGFP-G3BP1 while points outside this box and in the green and blue boxes correspond to higher order species. Panels **I-A, I-F,I-K** are control cells (n=8); Panels **I-B, I-G, I-L** are cells subjected to hypo-osmotic stress (150 mOsm, 5 min) (n=6); Panels **I-C, I-H, I-M** are cells subjected to arsenite stress (0.5mM, 10 min) (n=6); Panels **I-D, I-I, I-N** are cells subjected to carbachol stimulation (5μm, 10 min)(n=6); Panels **I-E, I-J, I-O** are cells subjected to heat shock (42°,60 min)(n=6); Scale bars are 10 μm long. Part ***II*** is similar to Part ***I*** for cells treated with siRNA(Cav1) (n=6) Scale bars are 10 μm long.

**Fig. 5B.II.**
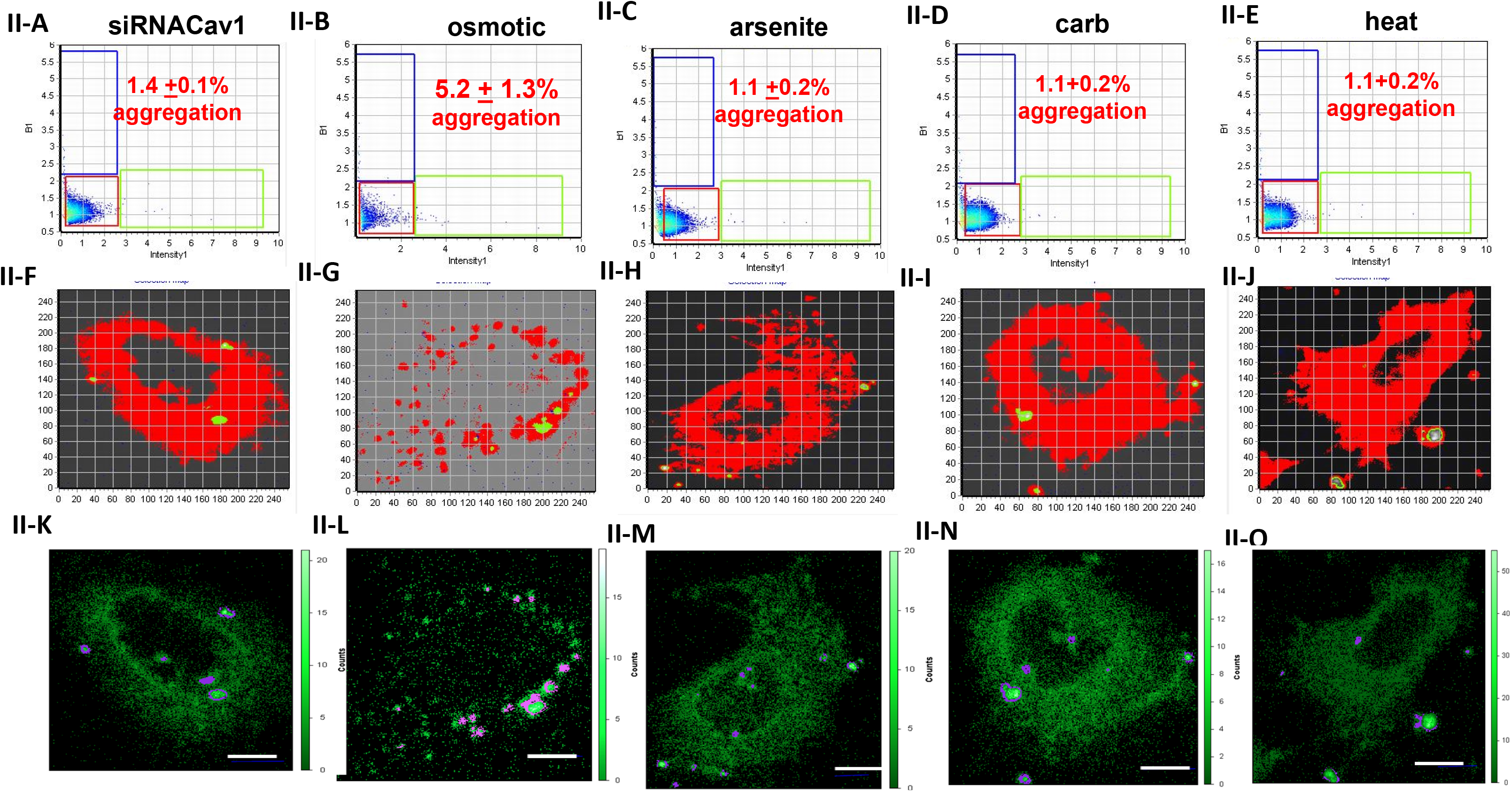
N&B analysis of eGFP-G3BP1 aggregation in Caveolin 1 downregulated WKO-3M22 cells.

It is also possible that reduced cytosolic RNA with Cav1/cavin-1 knock-down will impact p-bodies. We used the p-body marker, LSM14A, which is associated with inactive mRNA storage, to monitor the formation of p-bodies on the micron scale (Li et al., 2012; Parker and Sheth, 2007). High-resolution confocal images were analyzed to determine the number and area of LSM14A particles. Compared to control cells, down-regulating Cav1 had little effect on p-bodies, but down-regulating cavin-1 results in slightly larger LSM14A particles indicating increased p-body formation (**Fig. 6**). Additionally, we find that carbachol stimulation increases formation of LSM14A containing p-bodies while osmotic stress has no effect. Taken together, these studies suggest that lower RNA levels due to Cav1/Cavin knock-down do not affect p-bodies.

**Fig. 6.**
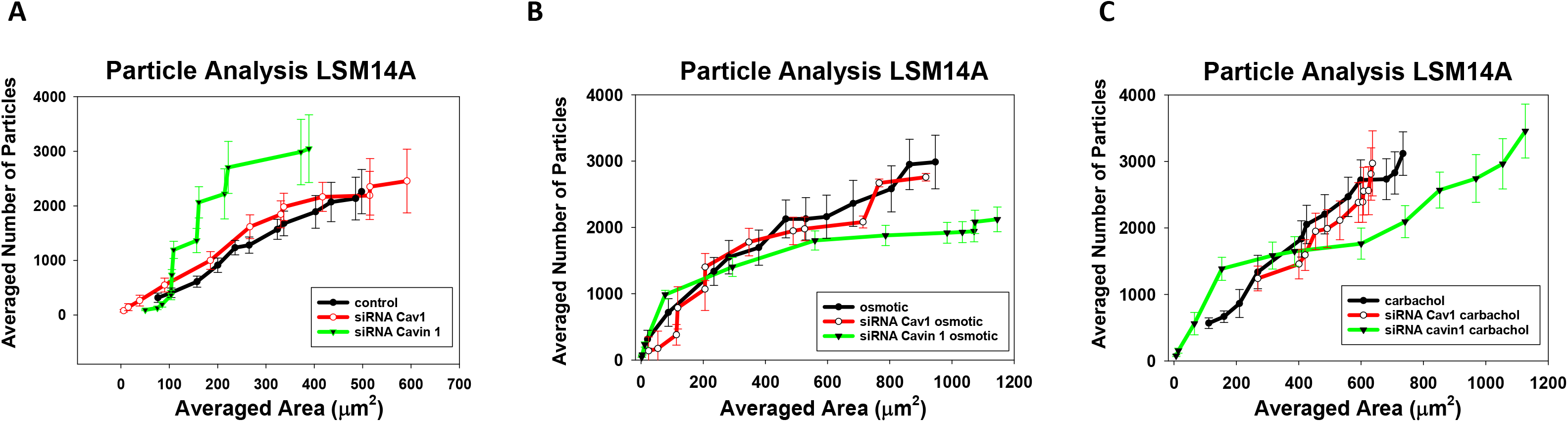
Cavin 1 impacts LSM14A containing P-body formation. The size and number of particles associated with monoclonal anti-LSM14A in the cytosol of fixed and immunostained WKO-3M22 cells was measured on a 100x objective and analyzed using Image J (see methods). Comparison of mock treated and cells treated with siRNA(cavin-1) or siRNA(Cav1) under basal conditions **(A),** hypo-osmotic stress (150 mOsm, 5 minutes) (**B),** and timulation of Gαq by treatment with 5 μM carbachol **(C).** All measurements are an average of 3 independent experiments that sampled 10 cells, where SD shown and where the p values was determined using ANOVA.

The studies above indicate that Cav1/cavin-1 levels primarily affect the formation of stress granules that form under various environmental conditions rather than p-bodies, which are associated with RNA storage and degradation under basal conditions. Therefore, we sought to investigate the effect of Cav1/cavin-1 on the formation of large and small aggregates containing Ago2, which aggregates in cells subjected to osmotic stress or Gαq activation (Qifti, Jackson et al., *in press*). We monitored stress-induced aggregation of eGFP-Ago2 in live cells with Cav1/cavin-1 down-regulation. When viewing Ago2 aggregates in control cells by fluorescence microscopy, we find that environmental stress does not significantly impact the size and number of Ago2 particles (**Table 1 and** *Supplemental 3*) and minor changes similar to PABPC1 particles are seen for cells treated with siRNA (Cav1). When we view small Ago2 particles in control cells by N&B analysis, we find a large impact by osmotic and carbachol stress but little change with heat and arsenite (**Table 1**). These results show that down-regulation of Cav1/cavin-1 reduces stress-induced aggregation, most likely due to reduced cytosolic RNA levels (*Supplemental 4*).

**Table 1:** Stress responses quantified by Particle Analysis and N&B. Legend – Shown are values for the slopes of the Particle Analysis curves (i.e. number versus area of particles), and the N&B values. Changes over four-fold are highlighted.

|  | <u>**Particle Analysis**</u> |  | <u>**N&B**</u> |  |
| --- | --- | --- | --- | --- |
|  | **Ago2** | **PABPC1** | **Ago2** | **G3BP1** |
| **Control** | 1.82 | 4.18 | n/a | n/a |
| **osmotic** | 1.63 | 5.56 | 34 | 7.6 |
| **carbachol** | 1.69 | 3.21 | 19 | 12.6 |
| **arsenite** | 1.55 | 2.87 | 5 | 3.9 |
| **heat** | 1.46 | 3.07 | 3.9 | 6.1 |
| **siRNA Cav1** | 2.47 | 7.62 | 5.3 | 1.4 |
| **siRNA Cav1 + osmotic** | 1.69 | 4.27 | 2.3 | 5.2 |
| **siRNA Cav1 + carbachol** | 3.12 | 4.43 | 0.4 | 1.1 |
| **siRNA Cav1 + arsenite** | 1.74 | 6.05 | 1.3 | 1.1 |
| **SiRNA Cav1 + heat** | 5.31 | 4.46 | 16.4 | 1.1 |
| **siRNA cavin-1** | 2.27 | 3.45 | 9.9 | 1.3 |
| **siRNA cavin-1 + osmotic** | 1.39 | 2.89 | 3.6 | 1.6 |
| **siRNA cavin-1 + carbachol** | 7.32 | 7.39 | 4 | 1.2 |
| **siRNA cavin-1 + arsenite** | 1.76 | 5.89 | 2.9 | 1.2 |
| **siRNA cavin-1 + heat** | 3.01 | 2.97 | 2.6 | 1.2 |

### Cavin-1 knock-out cells show widespread phenotypic changes as well as adaptive behavior

We obtained a strain of mouse embryonic fibroblasts (MEFs) where cavin-1 was knocked out using CRISPR/Cas9 genome editing technology (Liu and Pilch, 2016a). These knock-out cells have a threefold longer doubling time and a more trapezoid morphology as compared their wild type counterparts where the circularity dropped from 0.78±.2, n=16 to 0.55±.3, n=16. Additionally, we were unable to visualize caveolae by Cav1 immunostaining consistent with the absence of these caveolae, and these cells had a 55%-68% lower level of Cav-1 as quantified by western blot (*Supplemental 2B*). Surprisingly, we find that the cytosolic RNA levels of the knock-out cells were identical to the wild type in sharp contrast to transient down-regulation of Cav1/cavin-1.

### Cavin-1 knock-out cells show different cytosolic RNA size distributions

The adaptive changes in cytosolic RNA in the KO cells gives us the opportunity to understand the effects of cavin-1 at constant levels of cytosolic RNA. We find that even though the cytosolic RNA levels in the cavin-1 KO are the same as wild type, the size distribution are very different (**Fig. 7B-C**). Cytosolic RNA of wild type cells has two populations: a major one at small RNA sizes and a minor one at larger sizes. These populations are reversed in the knock-out cells; the peak at small RNA sizes was only 1/3 as high as the one at larger sizes. When we increase the expression of cavin-1 by over-expressing Cav1 in the knock-out cells, we find a shift in RNA sizes towards wild type values.

**Fig. 7.**
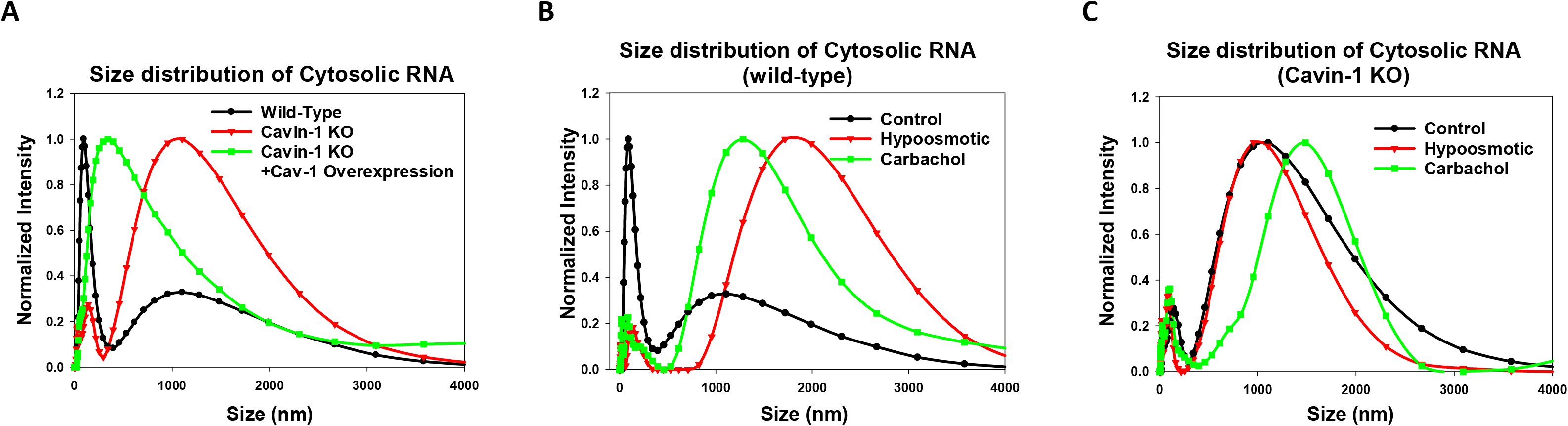
Differences in RNA size distribution between wild type and cavin-1 KO MEFs. **A-** Normalized DLS spectra showing changes in the size distribution of cytosolic RNA isolated from MEFs for mock treated control cells (black), cavin-1 knock-out cells (red) and cavin-1 KO cells over-expressing Cav1 (green). **B-** DLS spectra of cytosolic RNA from wild type MEFs under control conditions (black), hypo-osmotic stress (150 mOsm 5 min, red), cells over-expressing of constitutively active Gαq (green). **C-** An identical study showing the size distribution of cavin1 KO cells under control conditions (black), hypo-osmotic stress (150 mOsm 5min) (red) and Gαq stimulation (5 μM for 10 minutes, green). Each sample was scanned 3 times with 10 minutes per run. The number of independent samples were 6 per condition.

The shift to larger cytosolic RNAs sizes in the knock-out cells could be due to reduced processing. To test this idea, we subjected wild type cells to carbachol or hypo-osmotic stress to halt mRNA processing. These stresses reduced the amount of small RNAs consistent with the halting of Ago2 activity due to stress granule formation. The loss of small cytosolic RNAs in the wild type cells results in a DLS spectrum similar to unstressed knock-out cells. In contrast, subjecting cavin-1 knock-out cells to stress did not significantly change the cytosolic RNA size distribution. Thus, we interpret the higher sizes in the knock-out cells as being caused by reduced RNA processing **(Fig 8).**

**Fig. 8.**
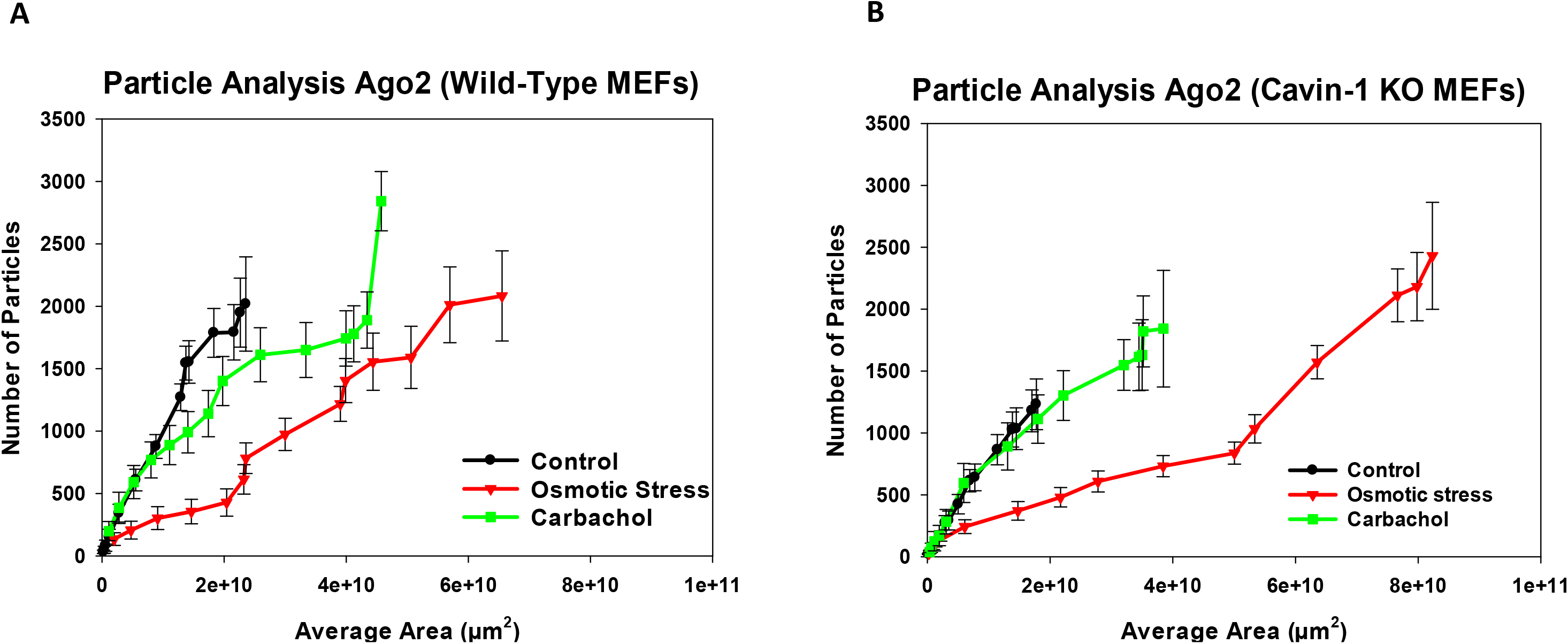
Formation of eGFP-Ago2 particles in MEFs cells under stress. Size and number of particles associated with anti-Ago2 in the cytosol of MEFs were measured on a 100x objective and analyzed using Image J (see methods). **A-** Treatment of wild type cells with hypo-osmotic stress (150 mOsm, 5 minutes-red) and Gαq stimulation (5 μM carbachol, 10 minutes – green). **B-** Treatment of cavin-1 KO cells to 5 minutes of osmotic stress (150 mOsm) and Gαq stimulation (5 μM carbachol, 10 minutes – green). Measurements are an average of 3 independent experiments that sampled 10 cells. P values were determined using ANOVA.

The large sizes of cytosolic RNA may be consistent with stalled ribosomal complexes (i.e., stress granules), and we determined the ability of the cavin-1 knock out cells to form stress granules. Particle analysis of wild type and KO cells showed Ago2 particles have are similar in size and number in cells under basal conditions and cells subjected to hypo-osmotic while the knock-out cells produced fewer particles under carbachol stimulation as compared to wild type cells (**Table 2,***Supplemental 5*). This reduction is consistent with the reduced levels of Gαq activation due to the loss of caveolae. The size of Ago2 particles formed were greatly enhanced upon Cav1/cavin-1 upregulation indicating some type of a compensatory mechanism (*Supplemental Table 2, S5*). These cells were also highly sensitive to arsenite suggesting that one of the adaptive mechanisms used by these cells is energy dependent. We also viewed small eGFP-Ago2 particles in wild type and cavin-1 knock-out cells by number and brightness (N&B) (*Supplemental Table 2*, *S5*). As compared to control cells, we find that the cavin-1 knock-out cells have eGFP-Ago2 particles that contain more fluorophores and are more intense. When cells are subjected to hypo-osmotic stress, the knock-out cells show enhanced aggregation which was slightly reversed with Cav1 overexpression (*Supplemental 6*). Thus, even though cavin-1 depleted cells have less particles on the micron scale, they have a larger number of smaller particles, and environmental stress promotes the formation of small Ago2 particles.

## DISCUSSION

In this work, we have begun to delineate the complex connections between caveolae, mechanical stretch and cell physiology. Caveolae provide mechanical strength to cells and serve as a platform that localizes signaling proteins (Parton, 2018). Several years ago, we found a connection between these two functions when we observed that Cav-1/-3 enhances Gαq / calcium signals, and that deformation of caveolae disrupts Cav/Gαq interactions (Yang and Scarlata, 2017). Here, we show that caveolae have an additional role by indirectly helping cells sense and respond to environmental stress conditions by modulating translational and transcriptional processes.

We began these studies noting that elevating cytosolic PLCβ reverses RNA-induced silencing of GAPDH but not Hsp90 though its inhibition of a promotor of RNA-silencing, C3PO (Philip et al., 2012), and that activation of Gαq drives PLCβ to the plasma membrane promoting both RNA-induced silencing and stress granule formation (Qifti et al., 2019b). Because Cav-1/-3 stabilizes activated Gαq (Sengupta et al., 2008), we postulated that its presence might also affect the production of these proteins by modulating Gαq activity. Our initial studies measured the effect of reduced Cav1 on the expression of two housekeeping proteins GAPDH and Hsp90, and the growth-associated protein Ras. As detailed below, the impact of Cav-1 on the basal levels of these proteins was difficult to quantify because of its broad effect on cellular RNA, and so we focused on changes in the levels of these proteins when caveolae deform under mild osmotic stress. By comparing to control cells, we find that levels of all three proteins are influenced by Cav1 down-regulation. The impact of Cav1 on GAPDH and Hsp90 does not appear to involve the Gαq/PLCβ signaling pathway since cells where either Gαq or PLCβ1 were down-regulated behaved similar to controls although Ras levels might be influenced by both Gαq and Cav1. Thus, these initial studies suggest that Cav1 may act through pathways distinct from Gαq/PLCβ to impact the levels of specific cellular proteins.

In searching for mechanisms that connect caveolae to protein levels, we noted that cavin-1 plays a dual role in cells by stabilizing caveolae domains on the plasma membrane and promoting the transcription of ribosomal RNA in the nucleus (Liu and Pilch, 2016b; Parton et al., 2018). Because expression of Cav1 and cavin-1 are linked, we were not surprised to find that down-regulating Cav-1 results in a dramatic decrease in cytosolic RNA levels, which may influence cellular levels of proteins like GAPDH, while down-regulating cavin-1 causes a much larger reduction of RNA.

Cavin-1 has been found to be released from the plasma membrane when caveolae domains disassemble, or upon phosphorylation during insulin signaling (Liu and Pilch, 2016a). Here, we worked at osmotic conditions where caveolae deform enough to disrupt Cav-1/Gαq interactions and focused on cavin-1 relocalization in real time in intact cells (Yang and Scarlata, 2017). We find that even these milder conditions promoted release of cavin-1 from the plasma membrane to the cytosol and nucleus. Not only did osmotic stress allow for cavin-1 relocalization, but carbachol stimulation did as well suggesting that strengthening Cav1-Gαq interactions weakened contacts with cavin-1. We were surprised to find that arsenite treatment, which has pervasive effects in cells also promotes cavin-1 relocalization to the nucleus. This relocalization may be due to initiation of cellular mechanisms to alleviate cell toxicity, or due to disintegration of caveolae-cytoskeletal contacts (Echarri and Del Pozo, 2015; Head et al., 2006; Parton et al., 1994).

While cavin1’s functions on the plasma membrane and nucleus are known, its role in the cytosol has not yet been determined. It is possible that the cytosolic population acts as a reservoir for the other cellular compartments, or that it serves an unknown cellular function. It is notable that cavin-1 complexes with cavin-2 in the cytosol (Breen et al., 2012). More recent work found that cavin-3, which interacts with both cavin-1 and caveolin-1 can also be released from the plasma membrane upon caveolae disassembly, interacts with a large number of cytosolic proteins and some of these interactions may overlap with cavin-1 (McMahon et al., 2019; Mohan et al., 2015). Pertinent for this study is the argument that cytosolic cavin-1 does not directly mediate stress granule formation since we have not detected cavin-1 in protein complexes associated with either PLCβ1 or Ago2 (Qifti et al., 2019b).

Our studies showed that reducing Cav1/cavin-1 levels and subjecting cells to osmotic stress reduced of GAPDH and Hsp90 but increased Ras relative to controls. We do not believe that these differential effects on transcripts are specific to cavin-1 but instead are indirect through other pathways, or due to the rate of translation of transcript caused by reduction ribosomes levels. We further explored this idea by assessing the relative size distribution of cytosolic RNAs. In WKO cells, lowering cavin-1 promotes the appearance of very large RNAs, while over-expressing cavin-1 increases small RNAs due to enhanced processing consistent with enhanced ribosomal activity. Reducing cytosolic PLCβ through stimulation of Gαq or application of osmotic stress shows the same shift towards smaller RNAs due to increased RISC activity while halting processing by arsenite causes a shift to very large sizes. These shifts are consistent with cavin-1’s role in promoting cytosolic RNA processing by increasing ribosomal RNA levels (Jansa et al., 1998).

Cavin-1 is highly expressed in proliferative tissues and ones rich in caveolae such as prostate cancer cells (PC-3), rhabdomyosarcoma (RMS), endothelial cells (ECs), etc. (Davalos et al., 2010; Faggi et al., 2015; Hill et al., 2008; Moon et al., 2014). Aside from making cells more susceptible to damage from mechanical stretch, down-regulation of cavin-1 would be expected to slow cell growth due to reduced ribosomal RNA output, and these characteristics were observed in the MEF cavin-1 knock-out cells (Liu and Pilch, 2016a). Even though they grew much slower than wild type, the KO cells had similar levels of cytosolic RNA showing that these cells have adapted to reduced ribosomal RNA transcription. We noted that the sizes of the RNAs in the cavin-1 KO cells were much larger than control suggesting a reducing in processing. When we over-express Cav1 in the KO cells, we find a shift in RNA sizes towards control cells. This shift is not due to increased expression of cavin-1 by Cav1 over-expression because the cavin-1 gene has been eliminated from these cells. One possibility is that Cav1’s stabilization of Gαq shifts the PLCβ population to the plasma membrane promoting RISC activity (Philip et al., 2012), and this idea is supported by previous FRET measurements (Qifti et al., 2019b).

Stress granule formation is initiated by cytosolic mRNA, and so it was not surprising that reducing Cav1/cavin1 results in cells that have impaired stress responses. Cavin-1 knock-out cells gave us to opportunity to assess the role of cavin-1 on stress granule formation at constant cytosolic RNA. We tested the importance of Cav1/cavin-1 on micron-sized particles imaging marker proteins, and on smaller aggregates by N&B analysis of fluorescent-tagged markers. Under basal conditions, the KO cells showed a greater number of small aggregates of Ago2 but a lower number of larger aggregates as compared to wild type cells and transfecting with Cav1 did not affect the results since cavin-1 is required for caveolae.

The overall goal of this study was to understand how caveolae transmits mechanical stress into cells. The mild stress used here was chosen to better emulate physiological conditions where caveolae are deformed but not completely disassembled (Yang and Scarlata, 2017). While we knew that this stress would reduce Gαq/calcium signals, we were surprised to find that caveolae changed the ability of cells to handle stress as seen by changes in protein content, and changes in cytosolic RNA, and changes in stress granule number and area. Our data indicate that these changes are mediated by two factors: relocalization of cavin-1 from the plasma membrane to the nucleus to allow increased rRNA production and increased translation, and increased RNA levels in the cytosol as described in the model shown in **Fig 9**. The nature of the specific proteins translated are unclear and may depend on competing transcripts, miRs and the rate of translation. Low levels of rRNA reduces the formation of ribosomes allowing cytosolic mRNA to be susceptible to inclusion in p-bodies or degradation by RISC (Eulalio et al., 2007). The rRNA resulting from nuclear cavin-1 provides a protective effect for cells by increasing cytosolic RNA levels allow for stress granule formation. Taken together, these studies show that cavin-1 can act as a sensor that communicates environmental stress to the nucleus that allow cells to better initiate stress responses.

**Fig. 9.**
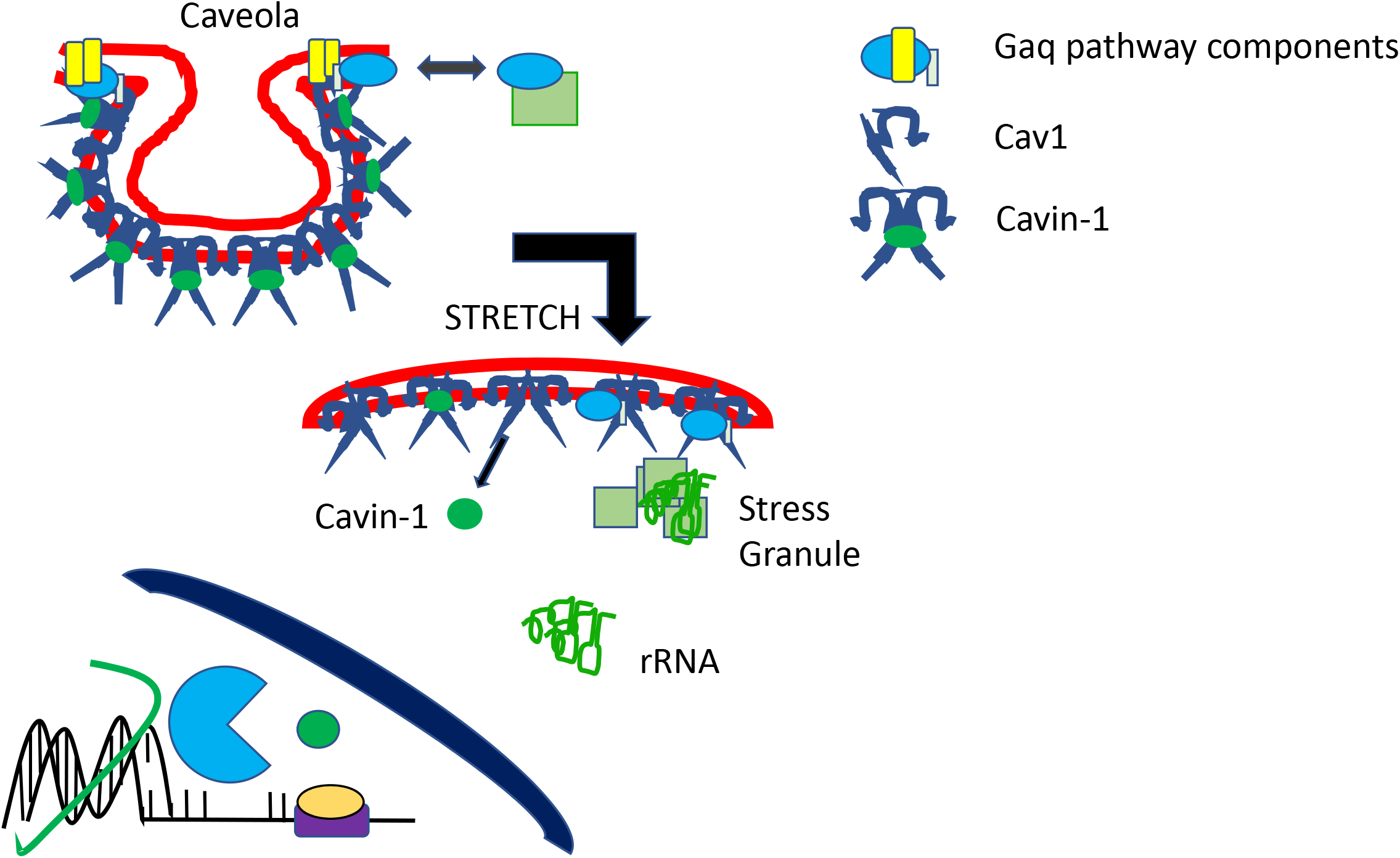
Model of stress effects and Cavin absence in cell structure. **A-** Gαq and PLCβ localize in caveolae; **B-** stretch flattens the domains releasing cavin-1 and shifting PLCβ to the cytosol reduce the amount of stress granules; **C** –cavin-1 relocalizes to the nucleus to influence transcription of specific genes that lead to changes in cell structure.

## MATERIALS AND METHODS

### Cell culture

Rat aortic smooth muscle (A10) cells were purchased from ATCC and used as described (Guo et al., 2015). Wystar Kyoto rat 3M22 (WKO-3M22) cells, originally obtained from ATCC, were a generous gift from Dr. Marsh Rolle. Mouse Embryonic Fibroplasts (MEFs) were a generous gift from Dr Liu Libin (Boston University School of Medicine). WKO-3M22 cell lines were cultured in high glucose DMEM (Corning) without L-glutamine with 10% fetal bovine serum, 1% sodium pyruvate, 1% non-essential amino acids (VWR) and 1% L-glutamine (VWR). MEFs were cultured in high glucose DMEM (Corning) without L-glutamine with 10% fetal bovine serum, 1% sodium pyruvate and 1% penicillin streptomycin. All cells were incubated at 37°C in 5% CO_2_.

### Plasmids & Stains

EGFP-hArgonaute-2 (eGFP-Ago2) was purchased from (Addgene plasmid # 21981) and was prepared in the laboratory of Philip Sharp (MIT). MCherry-Ago2 was a gift from Alissa Weaver (Vanderbilt University). EGFP-G3BP1 was purchased from Addgene (plasmid #119950) and was prepared in the laboratory of Jeffrey Chao (Friedrich Miescher Institute). EGFP- Cavin 1 was a generous gift of Dr Liu Libin (Boston University School of Medicine). CellMask Deep Red Plasma Membrane Stain was purchased by Thermofischer (cat# C10046) for the localization experiments. Plasmid transfections and siRNA knockdowns were done using Lipofectamine 3000 (Invitrogen) in antibiotic-free media. Medium was changed to one containing antibiotic (1% Penicillin/Streptomycin) 12-18 hours post transfection.

### Application of stress conditions

For the hypo-osmotic stress conditions, the medium was diluted with 50% water for 5 minutes before it was removed and replaced with Hank’s Balanced Salt Solution (HBSS) for imaging. For the arsenite treatment, a stock solution of 100mM arsenite in water was prepared. Cells were exposed to a final concentration of 0.5mM arsenite for 10 minutes before the medium was removed and replaced by HBSS for imaging. For the heat shock, cells were incubated at 40°C for 1 hour whereas for the cold shock cells were incubated at 12°C for 1 hour. For the oxidative stress treatment, a stock solution of 1M CoCl_2_ was prepared and cells were exposed to a final concentration of 1mM CoCl_2_ for 12-16 hours (overnight) before the medium was removed.

### RNA Extraction and Dynamic light scattering (DLS)

DLS measurements were carried out on a Malvern Panalytical Zetasizer Nano ZS instrument. For these experiments, mRNA from WKO-3M22 cells was extracted following the instructions from the Qiagen Mini Kit (Cat #: 74104). Prior to mRNA extraction, cells were exposed to stress conditions. For these measurements, approximately 50μLof extracted mRNA in RNase free water was added in a Hellma Fluorescence Quartz Cuvette (QS-3.00mm). Each sample was run 3 times, 10 minutes per run.

### Quantification of cavin-1 cellular localization by fluorescence imaging

Cell localization of eGFP-cavin-1 was assessed by combining multiple images of cells whose compartments were identified with fluorescent markers to identify the pixel coordinates of the plasma membrane, cytosolic and nuclear compartments, and tracking the shifts in eGFP-cavin-1 fluorescence intensity distribution relative to these coordinates in real time when cells are subjected to various stress conditions. A Horizontal Line Profile (H-Line Profile) was generated with counts vs pixel X where each intensity pixel corresponded to a unique coordinate location in the cell. These unique coordinates were matched to the plasma membrane, cytosolic and nucleus compartments based on the location of DAPI and the CellMask plasma membrane stain and the relative location of the eGFP-cavin1 to these compartments were calculated..

### Immunostaining and Particle analysis

Cells were grown to ~75% confluency and exposed to stress conditions, then fixed with 3.7% formaldehyde, permeabilized using 0.2% Triton X-100 in PBS then blocked using 100% Fetal Bovine Serum (FBS). Cells were then stained with primary antibodies incubated for 2 hours at room temperature or overnight at 4℃, washed and treated with a fluorescent secondary antibody for 1 hour. Some of the primary antibodies that were used were anti-PABPC1 (Santa Cruz sc-32318), antiG3BP1 (Santa Cruz sc-81940) and anti-LSM14A ABclonal Cat NO: A16682). After another wash, the 35mm MatTek glass bottom culture dishes were then imaged on the ISS Alba FLIM 2-photon confocal microscope using a 100X/1.49 oil TIRF objective to microscopically count the number of particles formed under different conditions per μm2. For each condition, 10-12 cells were randomly selected, and z-stack measurements were taken (1.0 μ/frame). Analysis was performed using ImageJ where each measurement was thresholded before averaging the number of particles per frame per measurement. Statistical significance was calculated using SigmaPlot with either a Student’s t-test, Tukey Test or Anova on Racks.

### Number and Brightness (N&B) measurements

N&B theory and measurement has been fully described (see (Digman et al., 2008)). Experimentally, we collected ~100 cell images viewing either free eGFP (control) or eGFP-Ago2, at a 66nm/pixel resolution and at a rate of 4 μs/pixel. Regions of interest (256×256 box) were analyzed from a 320×320 pixel image. Offset and noise were determined from the histograms of the dark counts performed every two measurements. Number and Brightness (N&B) data was analyzed using SimFC (www.lfd.uci.edu).

### N&B analysis

N&B defines the number of photons associated with a diffusing species by analyzing the variation of the fluorescence intensity in each pixel in the cell image. In this analysis, the apparent brightness, B, in each pixel is defined as the ratio of the variance, σ, over the average fluorescence intensity <k>:

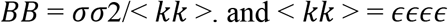

where n is the number of fluorophores. The determination of the variance in each pixel is obtained by rescanning the cell image for ~100 times as described above. The average fluorescence intensity, <k> is directly related to the molecular brightness, €, in units of photons per second per molecule, and n. B can also be expressed as

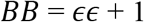

And the apparent number of molecules, N, as

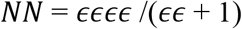

### Western blotting

Samples were placed in 6 well plates and collected in 250μL of lysis buffer that included NP-40 and protease inhibitors as mentioned before, sample buffer is added at 20% of the total volume. After SDS-PAGE electrophoresis, protein bands were transfer to nitrocellulose membrane (Bio-Rad, California USA). Primary antibodies include anti-PLCβ1 (Santa Cruz sc-5291), anti-Ago2 (abcam ab32381), anti-actin (abcam ab8226) and anti-eGFP (Santa Cruz sc-8334. Membranes were treated with antibodies diluted 1:1000 in 0.5% milk, washed 3 times for 5 minutes, before applying secondary antibiotic (anti-mouse or anti-rabbit from Santa Cruz) at a concentration of 1:2000. Membranes were washed 3 times for 10 minutes before imaging on a BioRad chemi-doc imager to determine the band intensities. Bands were measured at several sensitivities and exposure times to ensure the intensities were in a linear range. Data were analyzed using Image-J.

## ACKNOWEDGEMENTS

The authors are grateful to Liban Liu (Boston University School of Medicine) for providing wild type and cavin-1 knock-out MEFs and eGFP-cavin-1 DNA, and to Barbara Rosati (Stony Brook University) for sharing her RNA isolating procedure. The authors would also like to thank Siddartha Yerramilli, Katherine M. Pearce and Lela Jackson (Worcester Polytechnic Institute) for their helpful comments. This work was supported by NIH GM116178.

## Notes

### Competing Interest Statement

The authors have declared no competing interest.

